# B cell Compartmentalization in Blood and Cerebrospinal Fluid of HIV-Infected Ugandans with Cryptococcal Meningitis

**DOI:** 10.1101/759092

**Authors:** Samuel Okurut, David B. Meya, Freddie Bwanga, Joseph Olobo, Michael A. Eller, Fatim Cham-Jallow, Paul R. Bohjanen, Harsh Pratap, Brent E. Palmer, Katharine H. Hullsiek, Yukari C. Manabe, David R. Boulware, Edward N. Janoff

**Affiliations:** Research Department, Infectious Diseases Institute, Makerere University, Box 22418, Kampala, Uganda; Department of Microbiology, School of Biomedical Sciences, College of Health Sciences, Makerere University, 7072, Kampala, Uganda; Department of Medicine, School of Medicine, College of Health Sciences, Makerere University, 7072, Kampala, Uganda; Laboratory Department, Makerere University Walter Reed Project, 16524, Kampala, Uganda; Mucosal and Vaccine Research Program Colorado, Department of Medicine, University of Colorado Denver, Aurora, Colorado, USA; Denver Veterans Affairs Medical Center, Denver CO, 80045, USA; Division of Infectious Diseases and International Medicine, Department of Medicine, University of Minnesota, Minneapolis, MN, 55455, USA; U.S. Military HIV Research Program, Walter Reed Army Institute of Research, Silver Spring, MD, 20910, USA; Henry M. Jackson Foundation for the Advancement of Military Medicine, Bethesda, MD, 20817, USA; Division of Infectious Diseases, Department of Medicine, John Hopkins University School of Medicine, Baltimore, Maryland, MD, 21205, USA; Dept of Immunology and Molecular Biology, School of Biomedical Sciences, College of Health Sciences, Makerere University, Box 7072, Kampala, Uganda

**Keywords:** B cell subsets, activation, plasmablasts/plasma cells, PD-1, HIV, cryptococcal meningitis, survival

## Abstract

**Background:** Activated B cells modulate infection by differentiating into pathogen-specific antibody-producing effector plasmablasts/plasma cells, memory cells and immune regulatory B cells. In this context, the B cell phenotypes that infiltrate the central nervous system during HIV and cryptococcal meningitis co-infection are ill defined.

**Methods:** We characterized clinical parameters, mortality and B cell phenotypes in blood and CSF by flow cytometry in HIV-infected adults with cryptococcal (n=31), and non-cryptococcal meningitis (n=12), and heathy control subjects with neither infection (n=10).

**Results:** Activation of circulating B cells (CD21^low^) was significantly higher in blood of subjects with HIV infection compared with healthy controls, and greater yet in matched CSF B cells (p<0.001). Among B cell subsets, elevated frequencies of memory and plasmablasts/plasma cells most clearly distinguished the CSF from blood compartments. With cryptococcal meningitis, lower frequencies of expression of the regulatory protein PD-1 on plasmablasts/plasma cells in blood (median 7%) at presentation was associated with significantly decreased 28-day survival (29% (4/14 subjects)), whereas higher PD-1 expression (median 46%) characterized subjects with higher survival (88% (14/16 subjects)).

**Conclusion:** With HIV infection, B cell differentiation and regulatory markers are discrete elements of the circulating and CSF compartments with clinical implications for cryptococcal disease outcome, potentially due to their effects on the fungus and other local immune cells.

## Importance

Mortality from HIV associated cryptococcal meningitis co-infection remains abnormally high despite the use of optimal antifungal therapy and highly active antiretroviral therapy to treat HIV-1 and cryptococcal meningitis co-infected patients. However, it is not clear what contributes to the excess mortality after onset of cryptococcosis. We found an association of high percent expression of programmed death-1 (PD-1) receptor on plasmablasts/plasma cells population with host survival. We also found high expression of B cell activated cells and more mature B cell population in CSF compared to blood and this B cell dominant population in CSF of cryptococcosis patients expressed more PD-1. Together, these data in CSF and blood of cryptococcosis patients may informs further mechanistic studies of cryptococcus and host interaction to advance our understating of the possible pathways that may be targeted to influence pathogen control and host immune regulation to improve host survival.

## Introduction

Meningitis caused by the encapsulated fungus *Cryptococcus neoformans* is a leading cause of death among HIV-infected immune suppressed patients in sub-Saharan Africa, accounting for 15% of their deaths worldwide (1). Despite frequent exposure to the yeast in the environment, cryptococcal infection is very rare in healthy individuals. Immune status is a critical determinant of risk for fatal cryptococcosis. Type 1 helper T cells may contain primary infection in the lungs as a cornerstone of protection among healthy person (with ≈800-1200 circulating CD4+ T cells/μL) (2). However, most HIV-infected patients present with cryptococcal meningitis at a very advanced state of immunosuppression with CD4+ T cell counts <50 cells/µL (1).

B cells contribute to the development of a competent immune system by inducing naïve T cell activation, generating and maintaining serological memory, and regulating immune responses in health and in disease (3,4). In animal models, B cells produce antibodies against the cryptococcal polysaccharide capsule and other fungal antigens (5,6) that may attenuate infection and mediate fungal clearance (7). Specific antibodies may support opsonization and killing of the organism by phagocytes (8,9), neutralization of fungal virulence factors (10) or direct antibody-mediated toxicity and interference with fungal metabolism (7). B cells can produce either pro-inflammatory (e.g., IL-6, TNF-α and IFN-γ) (11) or anti-inflammatory cytokines (e.g., IL-10). IL-10-producing regulatory B cells, including plasma cells, modulate the activity of other immune cells in the local environment (4), as may B cells expressing surface immunomodulatory molecules such as PD-1 (12,13).

The contribution of pathogen-specific antifungal responses can be compromised during HIV-1 infection by polyclonal B cell activation and attenuated humoral responses to primary and recall antigens (14). Both *Cryptococcus* and HIV may have profound influences on B cell activation and differentiation and their effector and regulatory roles in the central nervous system (CNS) where most fatal cryptococcal disease occurs (15). To elucidate B cell signatures in AIDS-related cryptococcosis, we determined B cell phenotypes, activation and differentiation in blood and in CSF among persons with HIV with cryptococcal and non-cryptococcal meningitis and among HIV-negative healthy control subjects with neither infection and the association of these variables with survival.

## Results

### Subjects and mortality in HIV-associated meningitis co-infections

Age and gender did not differ significantly among the 3 study groups (Table 1). Circulating CD4+ T cell numbers were low in all HIV-infected subjects tested. CSF protein levels were similar among those with cryptococcal and non-cryptococcal meningitis. Although the Glasgow coma score was abnormal in only a quarter of subjects with cryptococcosis (<15 points), the 28-day mortality was high.

**Table 1.** Baseline characteristics of HIV infected participants with cryptococcal meningitis, non-cryptococcal meningitis and healthy control subjects.

| Groups | Cryptococcal meningitis |  | Non-cryptococcal meningitis |  | Healthy controls |  |  |
| --- | --- | --- | --- | --- | --- | --- | --- |
| Summary statistics | N | Median [IQR] or N[%] | N | Median [IQR] or N [%] | N | Median [IQR] or N [%] | P value |
| N | 31 |  | 12 |  | 10 |  |  |
| **Clinical** |  |  |  |  |  |  |  |
| Age [years] | 31 | 38 [32-41] | 12 | 39 [35-45] | 8 | 39 [36.5-46.5] | 0.81 |
| Female Gender | 31 | 8 [25.8%] | 12 | 6[50%] | 8 | 4[50.0%] | 0.13 |
| Weight [Kg] | 15 | 54 [50-58.0] | 3 | 60 [51.0-74.0] |  |  | 0.31 |
| CD4+ T cells/ul [normal 800-1200] | 22 | 13.5 [6-46.3] | 7 | 53.0 [7.0-279.0] |  |  | 0.3 |
| Currently on ART | 31 | 2 [6.5%] | 12 | 0 [0.0%] |  |  |  |
| Currently receiving TB therapy | 31 | 1 [3.2%] | 12 | 1 [8.3%] |  |  | 0.48 |
| Glasgow coma score <15 | 30 | 7 [23.3%] | 12 | 7 [58.3%] |  |  | 0.03 |

| CSF Parameters |  |  |  |  |  |
| --- | --- | --- | --- | --- | --- |
| Quantitative | 30 | 4.6 | 12 | 0[0-0] |  |
| cryptococcal |  | [4.1-5.3] |  |  |  |
| CFU/mL[Log10] |  |  |  |  |  |
| Sterile | 30 | 1[3.3%] | 12 | 12[100.0%] |  |
| cryptococcal |  |  |  |  |  |
| cultures |  |  |  |  |  |
| Opening pressure | 25 | 259 | 11 | 230 | 0.32 |
| [mm/H2O] |  | [160-370] |  | [128-278] |  |
| Opening pressure | 25 | 12[48.0%] | 11 | 3[27.3%] | 0.25 |
| >250 mm/H2O |  |  |  |  |  |
| Total WBC counts | 28 | 17.5 | 11 | 65 | 0.45 |
| [cells/μL] | | [4.0-112.5] | | [4.0-320.0] | |
| Total WBC counts | 28 | 15[53.6%] | 11 | 7[63.6%] | 0.57 |
| >5 [cells/μL] | | | | | |
| CSF protein | 30 | 59 | 12 | 95.5 | 0.16 |
| [mg/dL] |  | [20.0-97.0] |  | [43-187.0] |  |
| 1 Chi-square test for proportions. Kruskal-Wallis tests for continuous measures |  |  |  |  |  |
| 2 Kruskal-Wallis test for medians for comparing groups** |  |  |  |  |  |

WBC – White blood cells, CFU – colony forming units, PD-1 – Programmed death-1 inhibitory receptor, HIV+CM+ - HIV-infected and cryptococcal meningitis patients, HIV+ Non-CM: HIV-infected non-cryptococcal meningitis patients; Controls: healthy controls without HIV or without cryptococcosis. IQR – interquartile range.

Among all HIV-infected subjects with known outcomes, 42.9% (15/35) died in this period, including 40% (12/30; deaths) of those with cryptococcal meningitis. Among subjects with cryptococcal meningitis, median survival time was 10 days (95% Confidence interval (CI), 4-19 days) for those dying by 28-days and 50 days (95% CI, 32-99 days) for those dying after 28 days. Median survival was 19 days (95% CI, 9-30 days) for 4 subjects with *Mycobacterium tuberculosis* meningitis. One subject with meningitis of unknown cause died in 19 days.

### Overall B cell frequency and activation in blood and CSF among subjects with cryptococcosis

CD19+ B cells represented a greater proportion of circulating lymphocytes in blood among HIV-infected subjects with low CD4+ T cells compared with healthy controls, (medians, 12% in cryptococcosis, 27% in non-cryptococcosis and 4% in healthy controls; ANOVA, p<0.001) (Figure 2A). With HIV infection, B cells represented a higher proportion of lymphocytes in blood vs. CSF (medians, 12% vs. 2.3%, respectively; p<0.001) among cryptococcosis subjects and among non-cryptococcosis subjects (medians, 27% vs. 2.6%, respectively; p=0.011).

**Figure 1.**
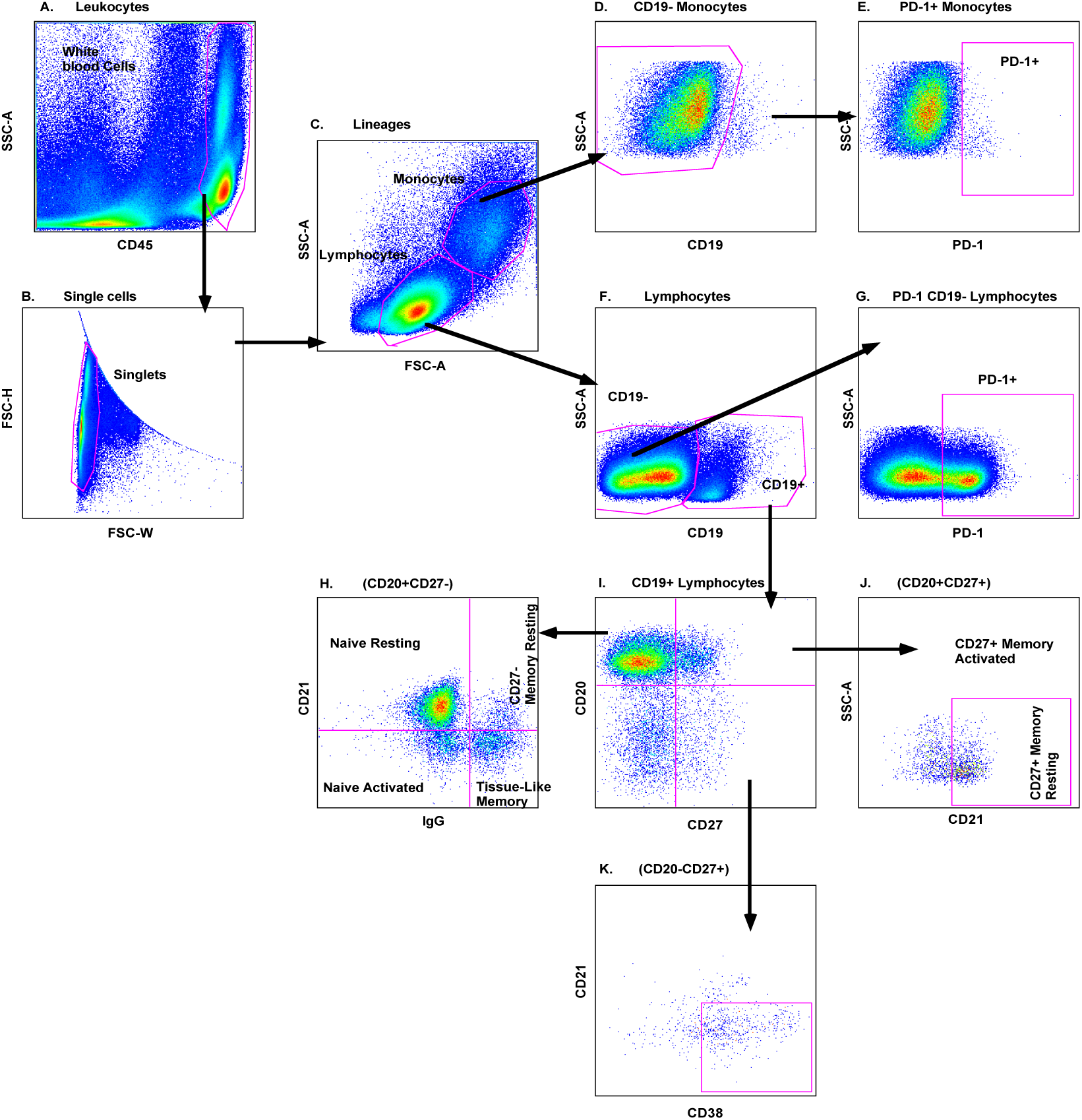
B cell gating strategy used to characterize activation and cellular differentiation in blood and CSF illustrated entirely on blood. Leukocytes are distinguished by expression of CD45+ expression [A] to exclude cryptococcal yeasts and selected for single lymphocytes [B-C]. PD-1 is gated on CD19-monocytes (D-E) and CD19-lymphocytes (F-G). B cell subsets (F-K) are defined per Supplementary Table 1, as described earlier (14,51). Gates indicated for PD-1, CD21 and CD38 were determined using fluorescence minus one cut-off [not shown], used to define subset expression, activation and PD-1 expression.

**Figure 2.**
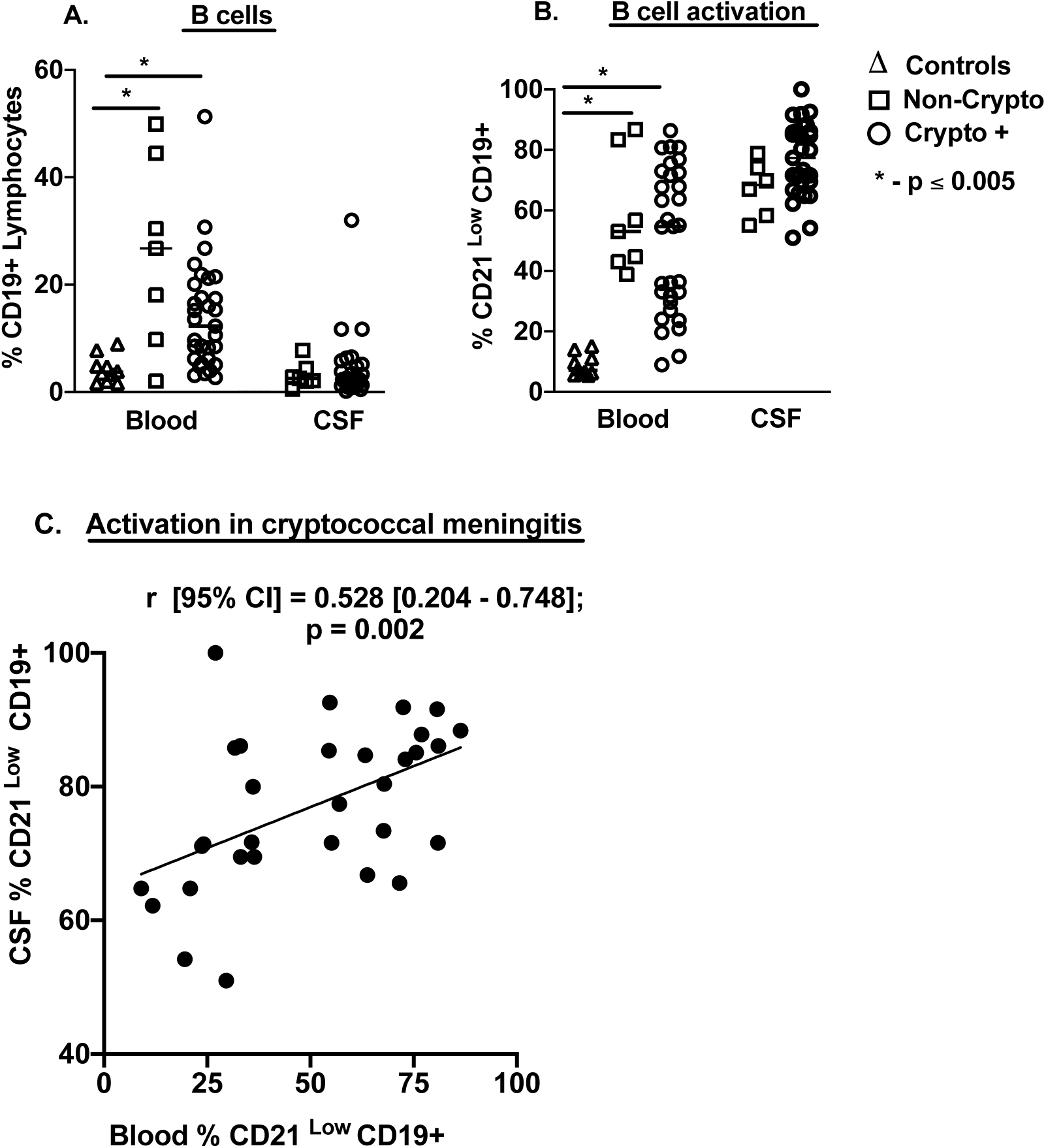
B cells Frequency and Activation in blood and in cerebrospinal fluid. Frequency of CD19+ B cells among lymphocytes **[A]**, and B cell Activation [CD21^lo^] by flow cytometry **[B-C]**. Samples were collected at presentation from Healthy controls subjects [blood samples; n=10], HIV-infected subjects with Cryptococcosis [blood n=31 and matched CSF n=31 samples] and with non-Cryptococcal meningitis [blood n=7 and CSF n=6 samples]. Values were compared using either Mann-Whitney U-test or using Kruskal Wallis and Spearman’s correlation coefficient. Horizontal bars indicate median values and * - asterisks show statistically significant results for p values <0.05.

B cell activation was significantly higher in both HIV-infected groups than among healthy controls in blood (medians, 55% and 53% vs. 7%, respectively, p<0.03) and higher yet in CSF, 68% and 77% (Figure 2B). Among cryptococcosis subjects, B cell activation in CSF positively correlated with that in blood (Figure 2C), but not among non-cryptococcal subjects (not shown).

### B cell subsets and activation in blood and CSF

Circulating B cells in blood showed distinct differences in subset distribution and activation. Naïve B cells predominated in blood in all groups (Figure 3A). Memory cells in blood were significantly lower among both HIV-infected groups compared with healthy control subjects. Tissue-like memory cells were over five-fold higher with HIV infection than in healthy controls. Plasmablasts/plasma cells, although a minority population in blood, were overrepresented with HIV infection (Figure 3A).

**Figure 3.**
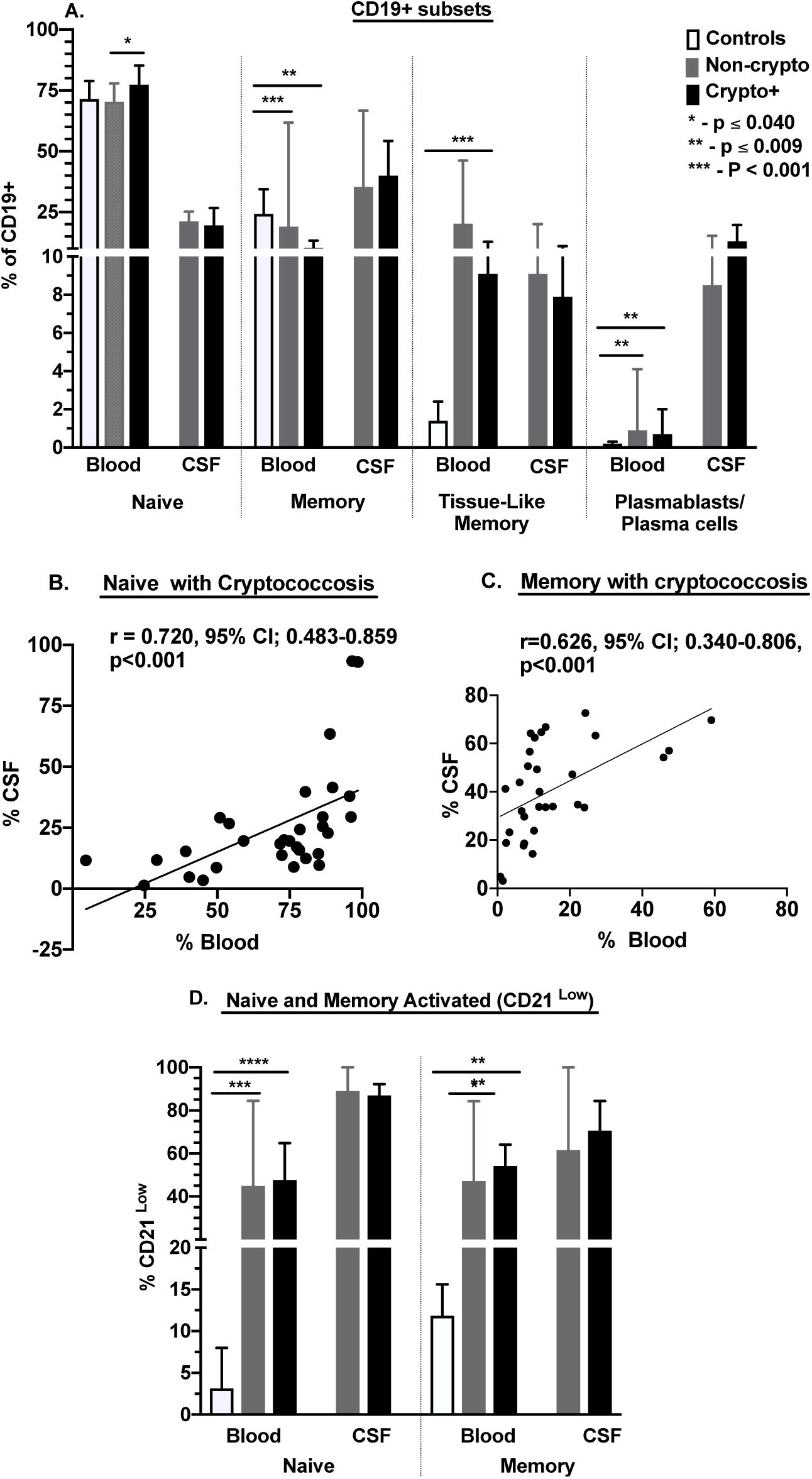
Distribution of B cell subsets and activation in blood and CSF. Frequency of B cell subsets among CD19+ lymphocytes by flow cytometry [defined in Supplemental Table 1]. Control samples [blood; n=10], cryptococcosis subjects samples [blood and CSF; n=31], and non-cryptococcosis subjects samples [blood & CSF; n=7]. Results are shown as medians [95% CI]. Values are compared using either Mann-Whitney U-test or Kruskal Wallis. * - asterisks show statistically significant results for p values <0.05.

In the CSF, B cells showed a more differentiated phenotype with naïve cells representing only about a quarter of cells compared with the majority in blood in all groups (Figure 3A); these proportions correlated in the two compartments (Figure 3B). Memory cells were also prominent in the CSF, accounting for up to half of B cells, and also correlated with those in blood (Figure 3C), suggesting trafficking between the two compartments. Plasmablasts/plasma cells frequencies in CSF greatly exceeded those in blood in HIV-infected subjects with (medians, 13% vs. 0.7%; p<0.001) and without (medians, 9% vs. 1%; p=0.008) cryptococcosis (Figure 3A).

In addition to subset differences between circulating B cells, patients with HIV demonstrated significantly higher levels of activation in both naïve and memory cells compared with healthy controls (Figure 3D). Activation was greater yet in B cells in the CSF, particularly among naïve cells, as well as among memory cells. These data indicate that B cells that traffic and localize to the CSF may be activated by infection at that site. That such activation is comparable in the presence or absence of *Cryptococcus* suggests that the local activating infection may be chronic HIV itself or the acute secondary pathogen. Thus, greater B cell differentiation characterizes the circulating B cell populations in HIV infection with or without cryptococcal meningitis infection, with prominent activated phenotypes being over expressed in the CSF.

### Preferential PD-1 expression on differentiated and activated B cells in blood and CSF

Programmed death-1 (PD-1) is a surface regulatory “check point” molecule identified prominently on T follicular helper cells and, less frequently, on B cells, NK cells, NKT cells and other myeloid derived cells (22). PD-1 was expressed on a minority of circulating B cells, but significantly more commonly on B cells in the CSF (Figure 4A). on CD19-lymphocytes, (majority T cells), PD-1 though most prominent in this cell population was more frequent in CSF than blood (Figure 4A). On the CD19-monocytes, was more frequently expressed on CSF of cryptococcosis patient than blood (Figure 4A). In blood, PD-1 expression was increased on more mature and on activated B cells, a pattern most directly applicable to healthy control subjects and those with cryptococcosis (Figure 4B). Most striking was the prominent high frequency of PD-1 on circulating CD27+ activated memory, on tissue-like memory and on plasmablasts/plasma cells in healthy controls and in the cryptococcosis co-infected group (Figure 4B).

**Figure 4.**
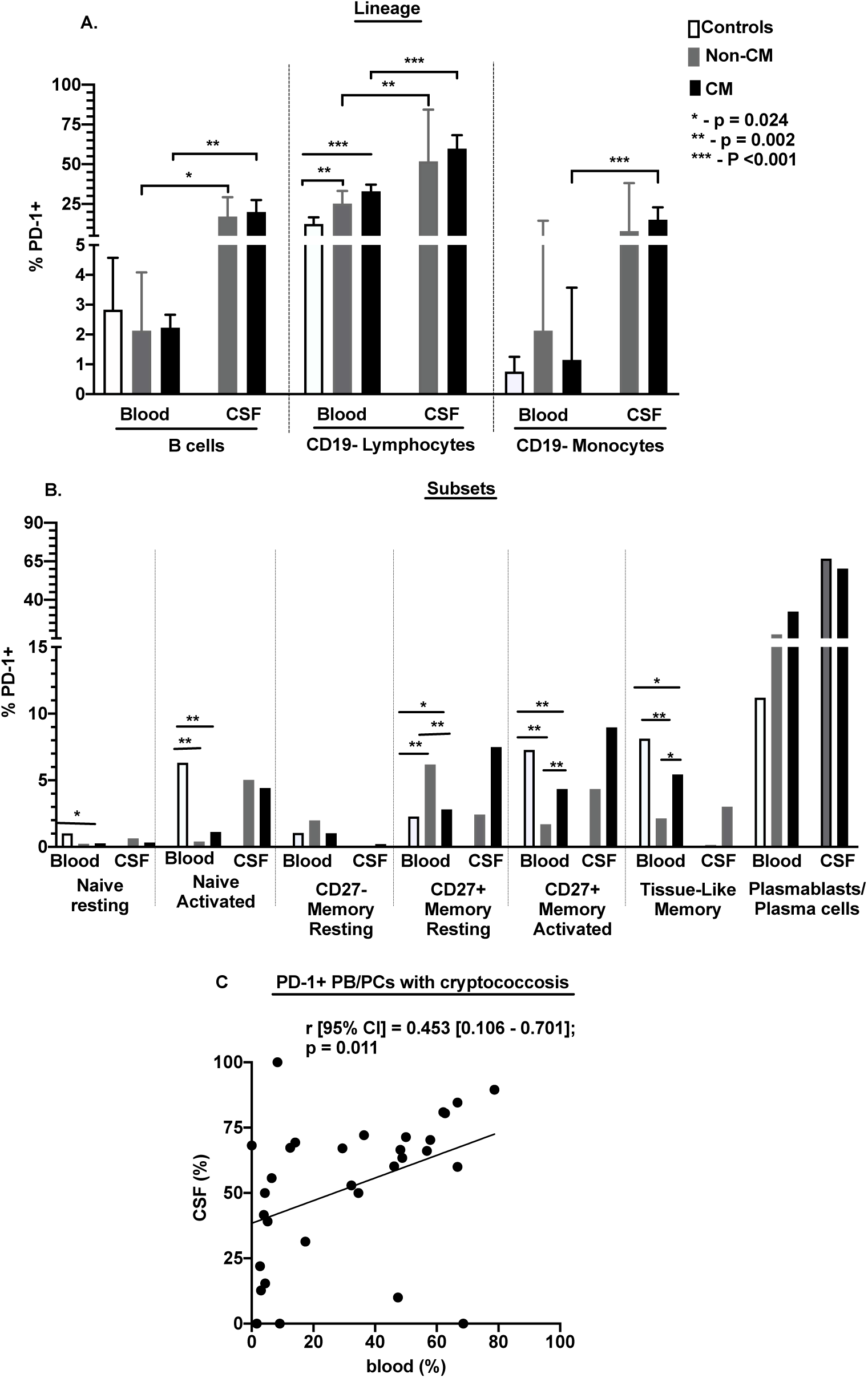
Expression of PD-1 on among healthy controls and among cryptococcosis subjects. Controls blood samples, [n=10], non-cryptococcosis samples, [blood; n=7 & CSF; n=7] and cryptococcosis samples, [blood; n=31 & CSF; n=31]. Results; **[A]**, PD-1+ expression measured as a frequency of CD19+ B cells **[B]**, Correlation of the frequency of PD-1+ expression on plasmablasts/plasma cells in blood and in CSF **[C]**, PD-1+ expression among B cell subsets. Bars show median values. Group values were compared using either Mann-Whitney U-test or using Kruskal Wallis. * = p value < 0.05.

Among other subsets, PD-1 was more commonly expressed on activated naive and CD27+ B cells from blood of healthy control subjects than of HIV-infected adults (Figure 4B and Supplementary Table S3). Among circulating B cells from adults with HIV infection, PD-1 was more prevalent on activated CD27+ memory activated and on Tissue-like memory, but not resting CD27+ memory B cells in those with *Cryptococcus* vs. non-cryptococcal meningitis.

As in blood, PD-1 in CSF was identified most frequently on PB/PC. These values were significantly associated in the two compartments among cryptococcosis subjects (Figure 4C). Thus, the preferential display of PD-1 on activated and differentiated memory B cells in the healthy controls and in those with cryptococcosis invokes the possibility of a regulatory role of this molecule on B cells in cryptococcal infection.

### PD-1+ expression on plasmablasts/plasma cells correlates with mortality among cryptococcal meningitis subjects

As noted above, overall mortality at 18 weeks was high in those with cryptococcal meningitis (18/30; 60%). Among the various B cell subsets, only PD-1+ expression on circulating plasmablasts/plasma cells was significantly associated with survival (also overall B cell activation by univariate but not multivariate analysis). Of the 10 subjects who died by 28 days after diagnosis with cryptococcosis, PD-1 was identified on a median of 7% of circulating plasmablasts/plasma cells. In sharp contrast, PD-1 was expressed by a median of 46% of these cells among 20 survivors (Figure 5B). Thus, low PD-1 expression was associated with early mortality and high PD-1 expression with survival.

**Figure 5.**
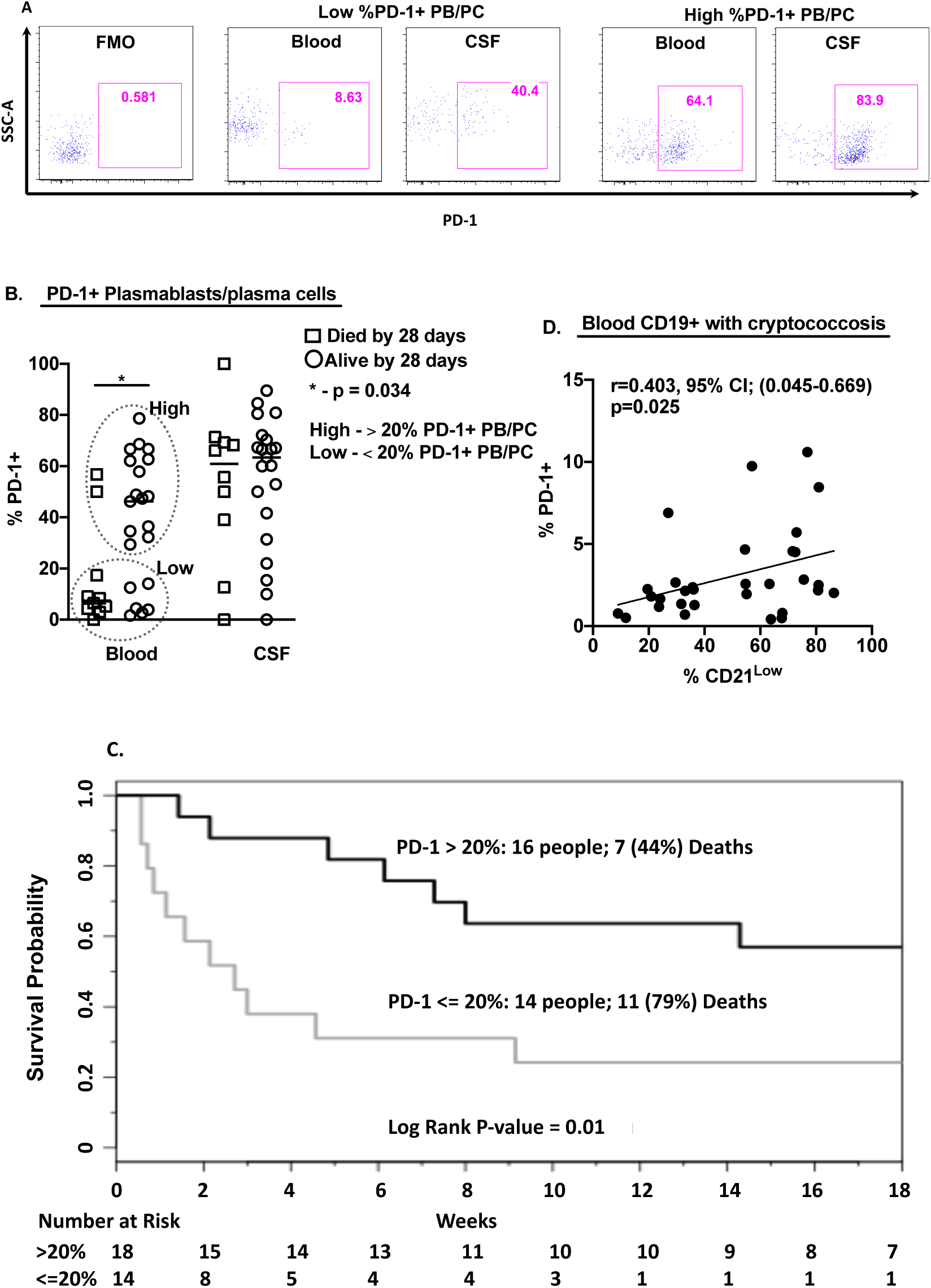
Programmed Death-1 expression on blood plasmablast/plasma cells at onset of cryptococcosis predicts 28-day survival or mortality. PD-1 – programmed death-1 receptor, PB/PC – Plasmablasts/plasma cells, Non-CM – HIV and non-cryptococcal meningitis co-infection, CM - Cryptococcal meningitis. **[A]** Individual profile of PD-1+ PB/PCs among low and high PD-1+ PB/PCs. **[B]** frequency of PD-1+ plasmablasts/plasma cells measured as frequency of PD-1+ expressing plasmablasts/plasma cells among cryptococcosis subjects who died by 28 days [n=10] Vs. Survivors at 28 days [n=20]. **[C]** Kaplan-Meier survival outcomes over time among cryptococcal meningitis subjects with low [<20%] [n=14] and high [>20%] [n=16] PD-1 expression on plasmablasts/plasma cells. **[D]** PD-1+ and CD21 low expression on B cells. Hazard ratio was determined as log rank of survival days. Censored subjects; Three cryptococcosis patients were lost to follow-up after hospital discharge and were excluded from the survival analysis. Mortality was determined at 18 weeks of follow-up.

Using PD-1+ plasmablasts/plasma cells as a continuous variable, every 5 units increase in PD-1+ expression on these cells was associated with 17% less chance of death in the acute setting by 28 days when most mortality occurred (HR (95% CI) = 0.83 (0.71, 0.98); p=0.02). This association of PD-1 on plasmablast/plasma cells at presentation was no longer significant with overall mortality at 140 days (18 weeks) (HR (95% CI) = 0.93 (0.84-1.02); p=0.13). In further exploration, using a cutoff point of 20% PD-1 plasmablasts/plasma cells expression, subjects with the PD-1 values ≤ 20% had had an increased risk of death (log rank p-value = 0.01; Figure 5C). Those with PD-1 values < 20% had a 7-fold increased risk of 28-day mortality compared to those with PD-1 > 20% (HR (95% CI) = 7.38 (1.69-34.36); p=0.01). While intriguing, the confidence interval is significant but wide, the p value is not very small, so this cutoff value is exploratory. By univariant analysis, only circulating B cell activation independently correlated with death (p=0.03); but not other reported risks for mortality (age, gender, Glasgow coma score, CSF protein, CSF white cell counts, or CSF fungal quantitative culture) (16,23). PD-1 expression on circulating B cells was associated with B cell activation in blood (Figure 5D). Whether these observations represent a plausible mechanistic impact of B cells and PD-1 on host survival or a secondary effect is under investigation.

## Discussion

The persistence of the very high mortality caused by cryptococcal meningitis during HIV infection in sub-Saharan Africa despite early diagnosis (24) drives efforts to improve antifungal therapy and complementary immune mechanisms of control. We describe for the first time, the distinct B cell subset maturation, activation and regulatory markers in paired CSF and blood from HIV-infected individuals with and without cryptococcal meningitis and associated early mortality. The recognition that cryptococcosis during advanced HIV disease is both a systemic and neurologic disease is supported by the consistently high levels of the antigen in both blood and CSF at the time of or preceding clinical diagnosis. The related B cell response to the infection in both compartments is supported by the correlation between B cell subsets, activation and PD-1 expression in blood and CSF identified herein.

Consistent with earlier reports, circulating B cells from untreated viremic HIV-infected patients show diffuse activation, deficits in memory B cell frequencies and increased representation of tissue-like memory and plasmablast/plasma cells (25–29). These results were consistent in patients with low CD4+ T cell numbers and meningitis due to *Cryptococcus* and other or indeterminate causes and substantially extend characterization of B cells in these co-infected subjects. We could not determine whether these prominent perturbations in blood B cell subsets and activation were due to the chronic effects of advanced HIV disease with an additional effect of acute secondary infection where cryptococcal antigen and symptoms develop 1-4 weeks prior to diagnosis (16,23,30,31). This distinction is important in determining whether patients with such advanced disease can actually generate a specific response to this systemic and local infection. That T cells to the fungus are detected in blood (32) and antibodies in blood (33,34) and CSF (Finn E, Okurut S, Janoff EN. manuscript in preparation) suggests that they can.

In this context, these data build on a limited but well-derived set of observations about the presence of B cells in the CSF during HIV infection. All studies have shown a predominance of T cells in CSF of HIV-infected subjects with and without *Cryptococcus* (21,35,36). Among HIV-1 infected adults without neurologic disease, uncharacterized B cells represented a small proportion (≈1%) of lymphocytes in the CSF, albeit more frequent than in heathy control subjects (35). We found that B cells in the CSF represented a median of 2.3% and 2.6% of lymphocytes with meningitis of *Cryptococcus* or other origin respectively. CSF B cells in our population were distinguished by prominent activation (median 77% and 68% CD21-B cells), a majority memory and tissue-like memory phenotype and high numbers of plasmablast/plasma cells (median 13 and 8% in the two meningitis groups). Akin to persons with multiple sclerosis (37,38), short-lived CSF plasmablasts were also reported to be increased in adults with HIV infection, particularly early in their course (39). That B cells and plasmablast frequencies in the CSF correlated with HIV RNA in the CSF and decreased with antiretroviral therapy in that study implicated the virus as a stimulus for the presence of these cells. B cells and short-lived plasma blasts were also increased with neurosyphilis and declined with therapy (40), highlighting that secondary infections can also elicit these cells. We could not distinguish between the more prominent plasmablasts and infrequent plasma cells described with our markers, but the increased frequency of B cells of memory and plasmablast/plasma cell with meningitis are consistent with the CNS responses to neurosyphilis, viral meningitis and in multiple sclerosis (37–42). Whether ectopic germinal centers are present in the brain with cryptococcal meningitis, which has a prominent component in the brain parenchyma, as were identified with neurosyphilis (40) as a source of local B cell generation for antibody production, is under investigation.

A distinctive feature of B cells in both blood and CSF in this study was the prominent expression of the check-point regulatory marker PD-1. PD-1 (CD279) is well-recognized as a co-inhibitory molecule, particularly during chronic viral infection, e.g., with HIV, when its expression is associated with CD8+ T cell exhaustion, low proliferative capacity and effector function and HIV disease progression (43). In addition to CD4+ and CD8+ T cells, PD-1 is also expressed upon activation of NK T cells, monocytes and B cells. As an immunomodulatory surface receptor, PD-1 on B cells can downregulate responses elicited through the antigen-specific B cell receptor (BCR) by dephosphorylating key cytoplasmic signal transducers of BCR signaling ((44).

PD-1 on CD4+ T follicular helper cells supports reversible inhibition of B cell responses via interactions with PD-L1 on B cells during HIV infection (23). However, PD-1 on B cells also has diverse and potent effects on B cell function with systemic impact. Among our subjects with cryptococcal and other forms of meningitis, PD-1 expression was increased on activated B cells, particularly CD27+ and tissue-like memory cells, as reported by others (43,45). Indeed, microbial antigens, modeled by toll-like receptor-9 agonists, augmented PD-1 expression on human B cells and IL-10 production (46). Akin to its effects on T cells, PD-1 on B cells can inhibit B cell activation, suppress B cell proliferation and impair B cell inflammatory cytokine responses (12,13,45,47).

Although less prominent on human B cells, PD-1 has been described at high levels among SIV-infected macaques in association with loss of memory B cells and fatal intestinal infection (48). The prominence of PD-1 on B cells in our patients with cryptococcal meningitis and the associated early and high mortality associated with its expression on plasma cells are consistent with these results in primates who also died with secondary infections. Blockade of PD-1 in the macaques augmented SIV-specific antibodies and improved survival. Thus, PD-1 on B cells may limit the antibody-producing effector function of B cells directly, and loss of innate-like IgM+ CD27+ memory B cells population that produces natural IgM antibodies is associated with risk of cryptococcosis among HIV infected individuals (49,50). PD-1 on B cells can also limit the effector activity of neighboring CD4+ and CD8+ T cells (14) and monocytes to clear cryptococcal infection by both PD-1-dependent and -independent IL-10-mediated regulatory mechanisms.

Thus, during cryptococcal infection in subjects with advanced HIV disease, PD-1 expression is prominently increased in frequency on B cells, and plasmablasts/plasma cells in particular, in both blood and CSF. The association of high PD-1 expression on circulating B cells with mortality suggests that this immunomodulatory protein may synergize B cell function, including antibody responses to *Cryptococcus*. Antibodies have been shown to facilitate control of his infection by mediating phagocytosis, effecting antibody dependent cytotoxicity by CD8+ T and NK cells and by directly inhibiting cryptococcal metabolism (7,8). Direct interactions of PD-1 with CD4+ and CD8+ T cells and monocytes, as well as production of IL-10, may also inhibit their ability to promote control of the organism in this population at high risk of serious disease complicated by neurologic sequelae and death. Control of chronic HIV infection will down-regulate PD-1 on T and B cells. However, mortality occurs early in the course of infection so, as shown in the SIV rhesus macaque model, blockade of PD-1 at this early stage may also promote both direct clearance and production of effective *Cryptococcus*-specific antibodies and enhance cellular immune function to promote clearance and limit relapse of this infection in persons with advanced HIV disease. We propose that understanding the mechanisms of induction of PD-1 on B cells in this setting and characterizing the consequent immune dysfunction will advance our understanding of the multiple facets of HIV-associated immune dysfunction and help prevent and resolve these serious opportunistic infections.

## Materials and Methods

### Study Participants

HIV-infected adults in Kampala, Uganda with cryptococcal meningitis (cryptococcosis) (n=31) and with non-cryptococcal meningitis; (non-cryptococcosis) (n=12) were selected from three prospective HIV meningitis clinical cohort studies: 1) Cryptococcal Optimal Antiretroviral Timing (COAT) trial (16), 2) Neurological Outcomes on Antiretroviral Therapy (NOAT) study (17) and 3) Adjunctive Sertraline for the Treatment of HIV Associated Cryptococcal Meningitis (ASTRO) trial (18). Inclusion criteria were; clinical evidence of meningitis, age ≥18 years, documented HIV infection, not receiving ART at enrollment and availability of cryopreserved cells from blood and CSF (Table 1). HIV infection was confirmed by bedside testing of previously undiagnosed subjects using the Ugandan Ministry of Health/WHO HIV testing algorithm. Cryptococcosis was confirmed by the presence of cryptococcal antigen by lateral flow assay (Immy Inc., Norman, Oklahoma, USA) in CSF and CSF cryptococcal quantitative cultures (19). The non-cryptococcal meningitis subjects where confirmed with tuberculosis and rifampicin gene xpert, 16S rRNA, and by quantitative fungal and bacterial culture as previously described in this HIV meningitis cohort (17). Of the non-cryptococcal meningitis co-infected subjects, 4 had *Mycobacterium tuberculosis* meningitis, 1 had neurosyphilis, toxoplasmosis and *Mycobacterium tuberculosis* co-infections while, 7 were without a known HIV co-infecting etiology. Whole blood, but not CSF, was obtained from healthy control subjects with neither HIV nor cryptococcal infection (n=10) from an HIV observational rural Ugandan cohort (20). Subjects prospectively provided informed written consent for the parent studies. Makerere University Research Ethics Committee granted ethical approval for use of stored specimens from previously consented adults. None of the subjects was on steroids, on ART or on anti-fungal therapy prior to sample collection.

### Sample Preparation

Peripheral blood mononuclear cells (blood) and CSF samples were collected within 72 hours of meningitis diagnosis. Blood and CSF cells were isolated and cryopreserved as previously described (21) in Roswell Park Memorial Institute enriched medium (69%) supplemented with 20% fetal bovine serum, 10% dimethyl sulphoxide and 1% penicillin/streptomycin in vapor phase in liquid nitrogen until testing. Blood and CSF cells were thawed, and cell viabilities and cell recoveries determined using an automated Guava PCA instrument before antibody staining for flow cytometry.

After thawing, median cell viability was 91% (range, 75-98%) from the CSF and 98% (Range 95–100%) from frozen PBMC. Median cells recovery was 1.2×10^6^ cells (Range 0.1-5×10^6^ million cells) per subject from CSF and 8×10^6^ (Range, 3-25×10^6^ cells) per subject from blood.

### Immunophenotyping

Thawed blood and CSF cells were stained with murine monoclonal antibodies reactive with human CD45 (FITC; clone HI30), CD20 (APC-Cy7; clone 2H7), and CD38 (PE-Cy7; clone HIT2) (Biolegend, San Diego, CA). CD19 (V500; clone HIB19) (BD Horizon, San Jose, CA); CD27 (PerCP-Cy5.5; clone M-T271), PD-1 (PE; clone EH12.1) and IgG (APC; clone G18-145) (BD Pharmingen, San Jose, CA) and CD21 (Pacific Blue; clone LT21; EXBIO Praha a. s., Czech Republic). CD45 expression was used to discriminate white blood cells in CSF and in blood from *Cryptococcus* yeast cells. Data were acquired using BD FACS Canto II, 8-color flow cytometer with BD Diva Software (BD Bioscience San Jose, CA, USA) and analyzed using FlowJo version 9.7.7 (Tree Star Ashland, Ore. USA). Gating was established for CD21, IgG, PD-1, CD38, CD20, and CD27 expression using fluorescence minus one controls. Spectral overlap was compensated using BD FACS compensation positive mouse Igκ beads and BD FACS Compensation negative mouse Igκ beads (BD Biosciences San Jose, CA, USA). A representative gating scheme is shown in Figure 1.

### B cell Differentiation and Activation

Based on results in Figure 1 and designations in Supplementary Table 1, CD45+CD19+ lymphocytes were characterized as resting or activated (CD21+ vs. CD21-respectively) naive (CD20+/CD27-/IgG-), memory (CD27+ or CD27-/CD20+IgG+CD38-), tissue-like memory (CD27-/CD21^low^/IgG+) B cells and plasmablasts/plasma cells (PB/PCs) (CD20-CD27++/CD38++/CD21^low^). Combined subsets accounted for 97.8-98.3% of gated CD19+ B cells in blood and 67.9% - 82.0% in the CSF in each group.

### Statistical Analysis

Data were analyzed using GraphPad Prism for Macintosh version 8.0 (San Diego, California, USA). Non-parametric Wilcoxon signed-rank test analyzed paired continuous variables, Mann-Whitney U-test and Kruskal Wallis tests analyzed unpaired continuous variables, and Kruskal Wallis test; analysis of variance (ANOVA) analyzed three group data. Survival was summarized with Kaplan-Meier plots and compared between groups with a log-rank test. Proportional hazards regression models were used to quantify the risk of death between groups. Survival data were censored at time of death, loss to follow-up, or at 18-weeks (the minimum follow-up time for the three studies). P-value ≤0.050 was considered statistically significant.

## Acknowledgements

We thank Joshua Rhein, Abdu Musubire, Nabeta Henry, Jane Francis Ndyetukira, Cynthia Ahimbisibwe, Florence Kugonza, Alisat Sadiq, Radha Rajasingham, Catherine Nanteza, Richard Kwizera, and Darlisha Williams from Infectious Diseases Institute for patient care. For flow cytometry research support, we thank Jeremy Rahkola, Tina M. Powell (Mucosal and Vaccine Research Center, University of Colorado Denver and Veterans Affairs Research Program) and Stefanie Sowinski and Olive Mbabazi (Infectious Diseases Institute, Makerere University Translational Sciences Laboratory). We thank Barbara Castelnuovo, Stephen Okoboi, Aidah Nanvuma, Angella Sandra Namwase from Infectious Diseases Institute, Research Department, and Research Capacity Development Unit for the mentorship and site supervision. We thank the Proscovia Naluyima, Alison Taylor, Britta Flach and the management of Makerere University Walter Reed Project Mulago Laboratory for supporting the project. We thank Damalie Nakanjako from the Department of Medicine, College of Health Sciences, Makerere University for healthy subjects specimens from her HIV seronegative observational cohort.

## Author contribution

Samuel Okurut, David B. Meya, Freddie Bwanga, Fatim Cham-Jallow, David R. Boulware, Edward N. Janoff, Yukari C. Manabe, Katharine H. Hullsiek and Michael A. Eller, conceptualized the study, contributed to framing of the research questions, participated in data analysis, drafting and revising the manuscript. Joseph Olobo, Paul R. Bohjanen, provided critical insights in manuscript revisions. Samuel Okurut, David B. Meya, Harsh Pratap and Brent Palmer designed the study panels, performed the immunological assays, analyzed the data and revised the manuscript.

## Figures

**Supplementary Table S1.** Markers of B cell subsets and activation.

| **B cell phenotypes** | **CD45** | **CD19** | **CD20** | **CD27** | **IgG** | **CD21** | **CD38** | **PD-1** |
| --- | --- | --- | --- | --- | --- | --- | --- | --- |
| **Naïve resting** | + | + | + | - | - | + | - | +/- |
| **Naïve activated** | + | + | + | - | - | - | - | +/- |
| **CD27- Memory** | + | + | + | - | + | + | - | +/- |
| **Resting** |  |  |  |  |  |  |  |  |
| **Tissue-Like** | + | + | + | - | + | - | - | +/- |
| **Memory** |  |  |  |  |  |  |  |  |
| **CD27+ Memory** | + | + | + | + | +/- | + | - | +/- |
| **resting** |  |  |  |  |  |  |  |  |
| **CD27+ Memory** | + | + | + | + | +/- | - | - | +/- |
| **activated** |  |  |  |  |  |  |  |  |
| **Plasmablast/** | + | + | - | + | +/- | - | + | +/- |
| **Plasma cells** |  |  |  |  |  |  |  |  |

Positive [+] shows presence of marker expression, negative [-] shows absence of marker expression and [+/-] show either presence or absence of marker expression.

**Supplementary Table S2.** B cell subsets and activation in cerebrospinal fluid [CSF] and blood among subjects with HIV-1 infection and cryptococcal and non-cryptococcal meningitis.

| Study group, | Blood | CSF | P value | Blood | CSF | P value |
| --- | --- | --- | --- | --- | --- | --- |
| B cell subsets | CM | CM |  | Non-CM | Non-CM |  |
| n | 31 | 31 |  | 7 | 7 |  |
| Naïve resting | 35.8<br>[18.9-60.5] | 2.6<br>[0.7-8.6] | <0.001 | 16<br>[10.9-39.9] | 2.6<br>[0.8-13.2] |  |
| Naïve activated | 28.4<br>[15.6-38.3] | 15.2<br>[8.5-22.8] | 0.007 | 53.1<br>[43.1-83.4] | 68.4<br>[57.5-75.3] |  |
| CD27- Memory resting | 1.4<br>[0.4-3.2] | 1.9<br>[0.3-4.5] |  | 2.3<br>[0.9-29.5] | 5.1<br>[2.9-11.9] |  |
| Tissue-like memory | 9.1<br>[3.4-20.2] | 7.9<br>[4.4-12.2] | 0.001 | 20.3<br>[6.5-22.4] | 9.1<br>[4.3-18.5] |  |
| CD27+ Memory resting | 3.5<br>[1.2-5.4] | 8.5<br>[3.4-13.9] | 0.001 | 5.2<br>[0.8-11.4] | 11.3<br>[8.7-22.4] |  |
| CD27+ Memory resting | 4.9<br>[2.0-9.3] | 27.2<br>[14.3-39.9] | <0.001 | 5.9<br>[2.7-17.7] | 24.7<br>[21.1-33.3] |  |
| Plasma | 0.7 | 13 | <0.001 | 0.9 | 8.5 | 0.008 |
| blasts/ | [0.2-2.4] | [3.3-20.9] |  | [0.6-1.3] | [4.4-14.2] |  |
| Plasma |  |  |  |  |  |  |
| cells |  |  |  |  |  |  |

All values as shown are median [IQR] percent of CD19+ B cells. P values were calculated with either Wilcoxon paired t-test; [Within cryptococcosis infected individuals: CSF vs. Blood comparisons] or Mann-Whitney unpaired U-test; [cryptococcosis vs. Non-cryptococcosis].

**Supplementary Table S3.** Programmed Death-1 [PD-1+] Expression on B Cell Subsets in CSF and in Blood with HIV infection with and without Cryptococcal Meningitis.

| **Study group,** | **Blood,** | **CSF,** | **P** | **Blood,** | **CSF, Non-** | **P** |
| --- | --- | --- | --- | --- | --- | --- |
| **PD-1+ B cells** | **CM** | **CM** | **value** | **Non-CM** | **CM** | **value** |
| **n** | **31** | **31** |  | **7** | **7** |  |
| **Naïve resting** | 0.3 | 0.3 |  | 0.2 | 0.6 |  |
|  | [0.2-1.0] | [0-2.6] |  | [0.1-2.0] | [0-2.2] |  |
| **Naïve activated** | 1.1 | 4.4 | **<0.001** | 0.4 | 5.0 |  |
|  | [0.7-1.9] | [1.4-7.7] |  | [0.2-1.3] | [1.3-16.5] |  |
| **CD27- Memory** | 1.0 | 0.2 |  | 1.9 | 0 | **0.009** |
| **resting** | [0.6-3.6] | [0-4.1] |  | [0-8.6] | [0-0.5] |  |
| **Tissue-like** | 5.4 | 3.0 | **0.003** | 2.2 | 0.1 |  |
| **memory** | [3.0-8.0] | [0.5-4.8] |  | [1.1-3.3] | [0-1.8] |  |
| **CD27+ Memory** | 2.8 | 7.5 | **0.001** | 6.2 | 2.4 |  |
| **resting** | [1.0-4.9] | [2.5-17.8] |  | [3.1-14.6] | [0-20.0] |  |
| **CD27+ Memory** | 4.4 | 9.0 | **0.001** | 1.7 | 4.4 |  |
| **activated** | [2.4-6.7] | [4.4-17.5] |  | [0.8-3.5] | [0-21.3] |  |
| **Plasmablasts/** | 32.3 | 60.2 | **<0.001** | 17.6 | 66.7 | **0.026** |
| **Plasma cells** | [5.2-56.8] | [31.4-70.3] |  | [5.9-26.1] | [44.5-100] |  |

All values as shown are medians [IQR] and percent of parent [B cells subsets]. P values were calculated with either Wilcoxon paired t-test; [within Cryptococcosis subjects; CSF Vs. Blood comparisons or Mann-Whitey unpaired U-test among non-cryptococcosis subjects Vs. Non-cryptococcosis subjects.

## Notes

Funding This research was supported in part by the National Institutes of Health (U01AI089244 (DRB, DBM), R01NS086312(DRB, DBM), K01TW010268 (DRB, DBM), R01AI108479 (ENJ)) and Veterans Affairs Research Service I01CX001464 (ENJ), the Fogarty International Center 1D43TW009771 (YCM), the GlaxoSmithKline-Trust in Science Africa (COL100044928) (SO). This work was in part supported by a cooperative agreement (W81XWH-07-2-0067) between the Henry M. Jackson Foundation for the Advancement of Military Medicine, Inc., and the U.S. Department of Defense (DOD). The views expressed are those of the authors and should not be construed to represent the positions of the U.S. Army or the Department of Defense. The funders had no role in the study design, data collection, data analysis, manuscript preparation and the decision to publish the work.

**Conflicts of Interest** Authors declare no conflict of interest and all the authors read and approved the final version of the manuscript.

